# Mouse pulmonary pathological characteristics induced by Asian-mineral dust transported with *Coniothyrium fuckelii* at a high altitude of 2,000 meters

**DOI:** 10.1101/2025.07.15.665011

**Authors:** Shotaro Yamano, Kaori Sadakane, Takamichi Ichinose, Yumi Umeda, Teruya Maki

## Abstract

Asian sand dust (ASD) events are known to transport not only mineral particles but also bioaerosols such as fungi across East Asia, posing complex respiratory health risks. Previously, we isolated *Coniothyrium fuckelii* (Strain Con-15H316)—a dematiaceous phytopathogenic fungus—from the ASD samples collected at an altitude of 2000 m over the Noto Peninsula, Japan. While Con-15H316 elicited potent immunological responses in our earlier cytokine-focused studies, its histopathological impact on mammalian lungs remained uncharacterized.

Here, we re-evaluated formalin-fixed murine lung tissues from mice intratracheally administered heat-treated ASD (H-ASD: only mineral particles), Con-15H316, and Con-15H316 in combination with H-ASD. H-ASD alone induced macrophage-dominant alveolar granulomas, and Con-15H316 alone triggered peribronchiolitis with eosinophilic infiltration. Strikingly, co-exposure resulted in a qualitatively distinct and synergistically worsened phenotype, featuring neutrophil-rich suppurative alveolar pneumonia, granulomas, and peribronchiolitis. Bronchoalveolar lavage fluid (BALF) analysis revealed marked co-elevation of lactate dehydrogenase (LDH) and CCL2 (C-C motif chemokine 2, also known as monocyte chemoattractant protein-1, MCP-1), suggesting concurrent tissue injury and monocyte-driven immune activation. To elucidate the cellular origin of the CCL2 protein, we reanalyzed two publicly available single-cell RNA-seq datasets, identifying monocytes and interstitial macrophages as the dominant Ccl2-expressing populations in microbially injured lungs.

Although *C. fuckelii* has been recognized generally as a plant pathogen, this study exhibited the first histological evidence of the potential of this fungus to induce or exacerbate mammalian lung injury, especially in combination with mineral dust. Our findings uncover a novel, non-allergic alveolar injury phenotype associated with microbial–particulate co-exposure, expanding current toxicological paradigms beyond pathologies associated with excessive immune reactions to inhaled particulates and bioaerosols.

## Introduction

Asian sand dust (ASD) provides several significant risks to human health^1^. The primary sources of ASD are the arid regions of northern China, particularly the Taklamakan and Gobi Deserts, from which large scale dust storms disperse ASD throughout East Asia^2^. ASD is known to carry a substantial load of bioaerosols, including both bacteria and fungi, into the atmosphere^3^. Surveys of the effects of dust storms in northern China revealed that dust storms resulted in a dramatic increase in the diversity and quantity of airborne bacteria in downwind areas such as the Korean Peninsula and Japan^4,5^. The airborne microbial cells associated with these dust events in Asia are known to heighten mortality rates and increase the incidence of respiratory and cardiovascular diseases^6^, as well as cause a rise in asthma cases^7^. Furthermore, co-exposure to dust particulates and microorganisms has been shown to exacerbate symptoms and diminish lung function of asthma patients during dust events in South Korea^8^ and Japan^9^.

In our previous study, fungal fragments and hyphae were collected successfully from ASD aerosols at high altitude using balloons and helicopters during ASD events in the Noto Peninsula, Japan^10^. Five fungal species were identified in isolates from the ASD aerosols: *Bjerkandera adusta* (Bje-07B507), *Lecythophora* sp. (Lec-13H319), *Cladosporium cladosporioides* (Cla-16H615), *Phialocephala sphaeroides* (Phi-15H321), and *Coniothyrium fuckelii* (Con-15H316). We assessed their health implications by administering fungal cells inactivated with 1% formalin, heat-treated ASD (H-ASD: only mineral particles), and their mixtures into the lungs of BALB/c mice. This study confirmed the serious respiratory toxicity of fungi debris in ASD particles. Notably, Con-15H316 associated with H-ASD exhibited the most pronounced pulmonary toxicity. This was characterized by elevated levels of various inflammatory cytokines in the bronchoalveolar lavage fluid and eosinophil infiltration in the bronchial interstitium. However, a histopathological examination of the lung tissues from these mice has yet to be fully conducted.

Additionally, we have previously performed extensive large-scale animal studies in accordance with OECD test guidelines^12^, using rodents and a whole-body inhalation exposure system, which have pathologically identified various lung diseases induced by nanomaterials such as multi-walled carbon nanotubes (MWCNT)^13–15^ and titanium dioxide nanoparticles (TiO_2_ NP)^16–19^, as well as organic dust like cross-linked water-soluble acrylic acid polymer (CWAAP)^20,21^ and gaseous substances such as 2-bromopropane^22^. Based on these results, we have acquired comprehensive insights into the histopathological diversity of aerosol-induced lung diseases in rodents.

In this study, we prepared lung tissue sections from archived paraffin-embedded blocks obtained in our previous study^10^ and re-analyzed previously reported bronchoalveolar lavage fluid (BALF) data to clarify additional histopathological features induced by H-ASD alone, Con-15H316 alone, and the combination of H-ASD and Con-15H316.

## Results

### Representative Histopathological Findings in H-ASD-Exposed Mouse Lungs

Histopathological analysis of the mouse lungs exposed to H-ASD alone at low magnification indicated no overt abnormalities (Figs. 1A and 1B). However, under polarized light at higher magnification, birefringent yellow dust particles—consistent with H-ASD—were occasionally observed within the alveolar spaces (Figs. 1C and 1D). Most particles were phagocytosed by alveolar macrophages, often forming small granulomatous foci in the alveolar regions (Fig. 1E). Notably, the interstitium surrounding bronchioles, blood vessels, and pleura remained histologically unremarkable and comparable to controls (Fig. S1). H-ASD particles were evenly distributed across all lung lobes, indicating successful and uniform intratracheal delivery. These findings indicate that H-ASD exposure alone induces granuloma formation and primarily localized to the alveolar regions.

**Figure 1.**
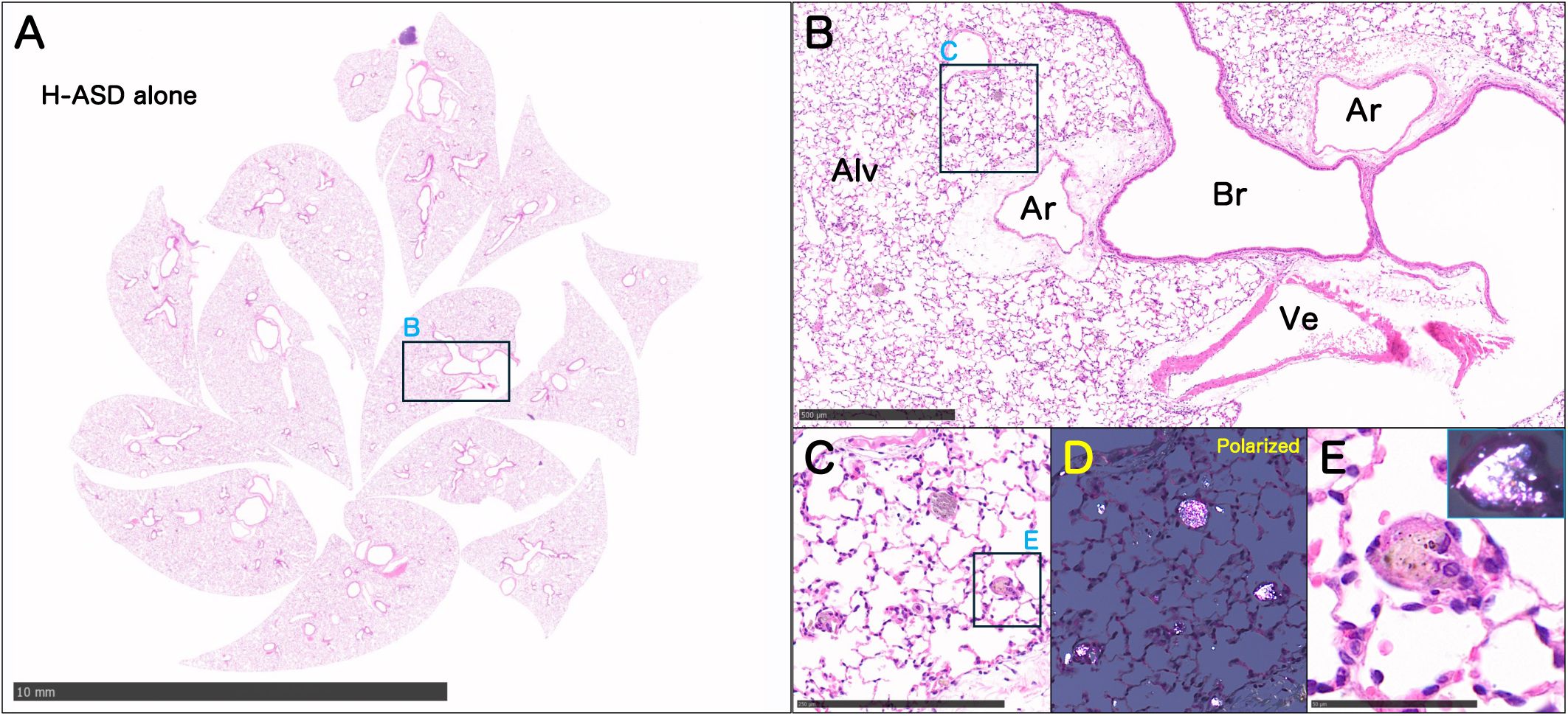
Histopathological findings of mouse lungs exposed to H-ASD alone. (A, B) Low-magnification H&E images reveal no overt histological abnormalities. (C, D) At higher magnification with polarized light microscopy, birefringent yellow H-ASD particles are observed within alveolar airspaces. (E) Granuloma formation is noted in alveolar regions, with most particles phagocytosed by macrophages. No significant peribronchial or vascular inflammation is present. Alv, alveoli; Br, bronchiole; Ve, vein; Ar, artery.

### Representative Histopathological Findings in Con-15H316-Exposed Mouse Lungs

Exposure to low amounts of Con-15H316 inactivated with 1% formalin resulted in patchy basophilic areas with increased cellularity, particularly around bronchi and blood vessels (Figs. 2A and 2B). Exposure to higher amounts of Con-15H316 increased the level of inflammatory cell infiltration in the peribronchovascular interstitium (Figs. 2C–2E), which was predominantly composed of lymphocytes and eosinophils. Mild interstitial collagen deposition was also noted in some regions (Fig. 2D). Goblet cell hyperplasia was evident in the bronchial epithelium, while the thickness of the bronchial smooth muscle remained unchanged. In the alveolar regions, patchy infiltrates of neutrophils, eosinophils, macrophages, and cellular debris were observed (Fig. 2F).

**Figure 2.**
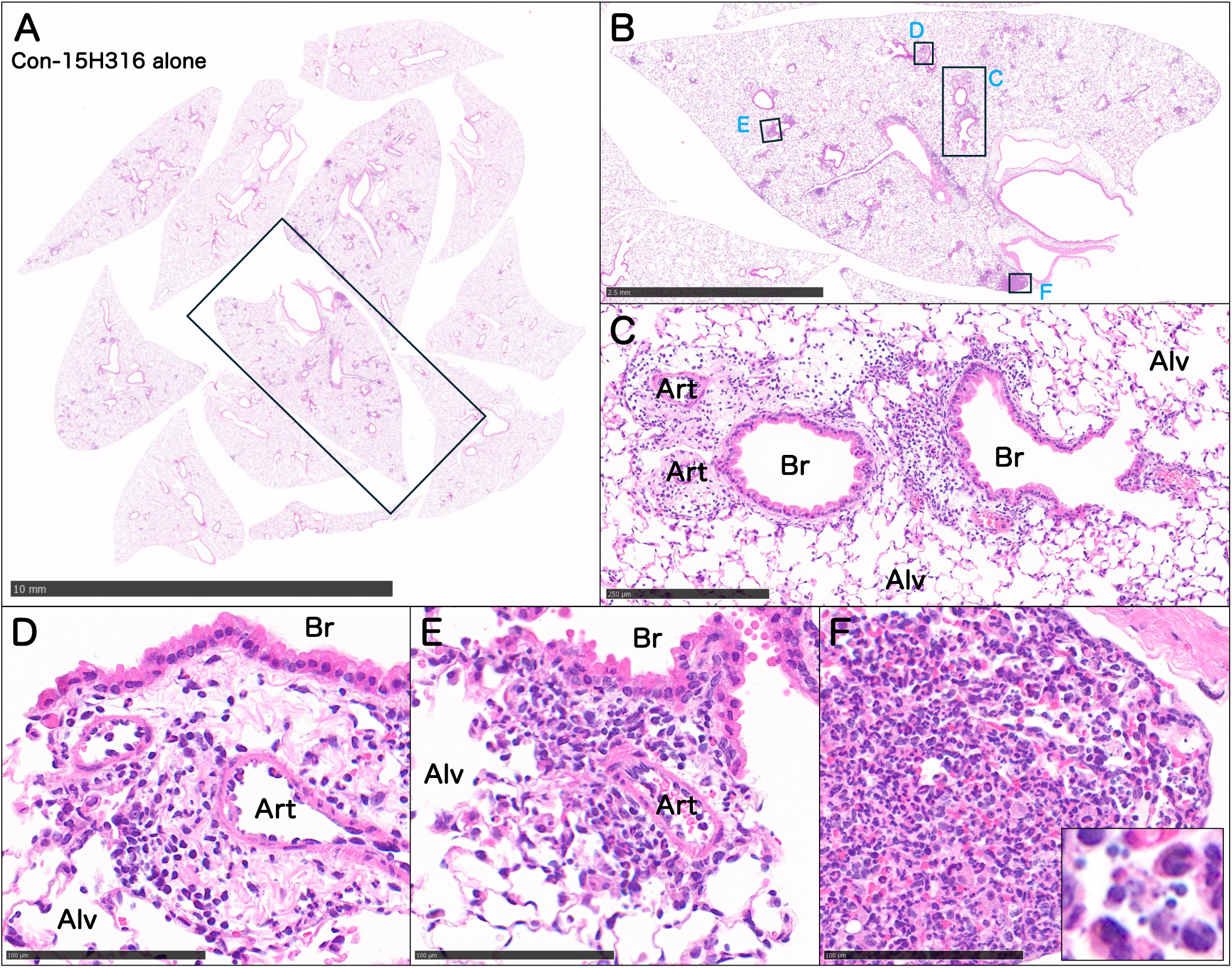
Histopathological findings in mouse lungs exposed to Coniothyrium fuckelii (Con-15H316) alone. (A, B) Patchy basophilic areas around bronchi and blood vessels suggest localized inflammation. (C–E) High-power images reveal dense eosinophilic and lymphocytic infiltration in the peribronchovascular interstitium, with mild collagen deposition. Goblet cell hyperplasia is present in bronchial epithelium. (F) Alveolar regions show patchy neutrophilic inflammation with cellular debris. Br, bronchiole; Art, arteriole; Alv, alveoli.

Occasional basophilic granules within these areas suggested the possible presence of fungal material. However, no granuloma formation or diffuse alveolar damage was observed in most regions. These results indicate that Con-15H316 exposure alone induced peribronchiolitis characterized by eosinophilic infiltration in the peribronchovascular stroma and goblet cell hyperplasia in the bronchial epithelium.

### Representative Histopathological Findings after Combined H-ASD and Con-15H316 Exposure

Mice exposed to a combination of H-ASD and Con-15H316 exhibited extensive and diffuse basophilic areas at low and intermediate magnification in comparison to exposure to H-ASD or Con-15H316 alone (Figs. 3A and 3B). When exposed to higher-amounts of H-ASD and Con-15H316, peribronchiolitis with eosinophilic infiltration and goblet cell hyperplasia was observed (Fig. 3C). These phenomena are similar to the effects of exposure to Con-15H316 alone, but the severity level in the mice exposed to the combination of H-ASD and Con-15H316 was much higher. Interestingly, neither free H-ASD particles nor particle-laden macrophages were observed within the peribronchovascular interstitium, suggesting that inflammation in this region was primarily driven by Con-15H316. In the alveolar regions, increased cellularity was evident (Figs. 3D–3E). High-power views revealed a complex inflammatory pattern composed of neutrophils, macrophages, and cellular debris (Fig. 3F). Multinucleated giant cells and H-ASD-laden macrophages formed nodular aggregates, consistent with granulomatous inflammation. In parallel, focal accumulations of neutrophils and nuclear debris indicated the presence of early suppurative pneumonia. Polarized light microscopy confirmed the retention of birefringent yellow H-ASD particles in affected alveolar spaces, indicating enhanced particle deposition under co-exposure conditions. While alveolar septal thickening was not evident, septal capillaries often showed increased intravascular leukocytes, suggestive of active immune cell recruitment.

**Figure 3.**
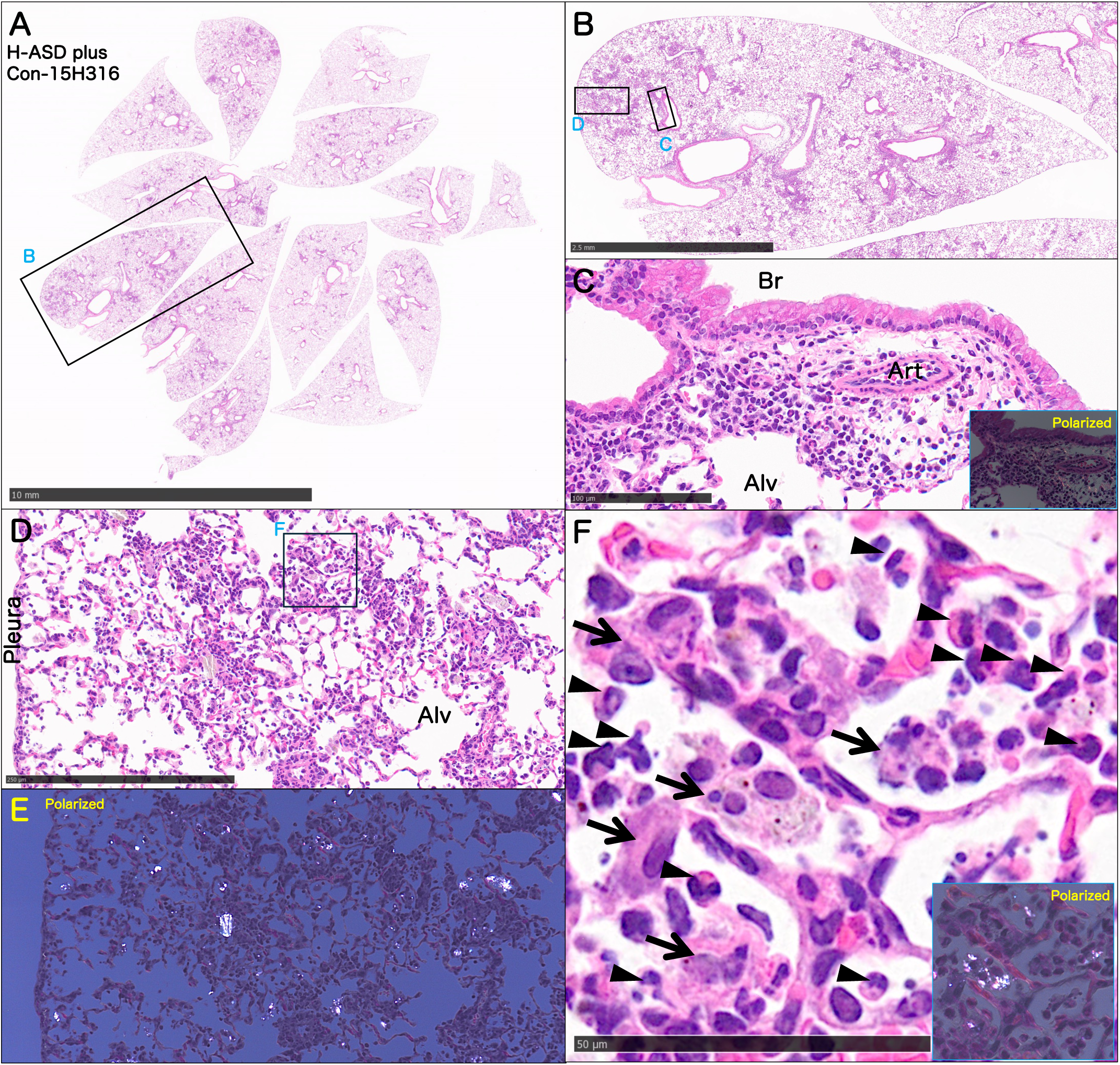
Histological findings in mouse lungs co-exposed to H-ASD and Con-15H316. (A, B) Extensive basophilic inflammatory lesions are observed at low and intermediate magnifications. (C) Severe bronchitis characterized by eosinophilic infiltration and cup cell hyperplasia. (D–F) Mixed inflammatory infiltrates composed of neutrophils, macrophages, lymphocytes, and multinucleated giant cells are observed in the alveolar regions. Polarized light microscopy (F) confirms increased accumulation of birefringent H-ASD particles in the alveolar spaces. In panel (F), the arrowheads indicate neutrophil clusters (pyogenic inflammation), and the arrows indicate macrophages phagocytosing H-ASD particles (granulomatous component), indicating that this alveolar region exhibits a hybrid inflammatory phenotype. Br, bronchus; Art, artery; Alv, alveoli.

Collectively, these findings indicate that co-exposure to H-ASD and Con-15H316 induces a hybrid inflammatory phenotype in the lung, combining particle-induced granulomatous inflammation and fungus-associated suppurative lesions, particularly within the alveolar compartments.

### Comparison of Histopathological Features and Bronchoalveolar Lavage Fluid (BALF) Markers Across Experimental Groups

Histopathological comparisons among the saline and three particle treated experimental groups focused on three lesion types: peribronchiolitis in the airway region, and granulomatous and suppurative inflammation in the alveolar regions (Figs. 4A–4C). The levels of the inflammation-associated biomarkers lactate dehydrogenase (LDH) and CCL2 (C-C motif chemokine 2, also known as monocyte chemoattractant protein-1, MCP-1) were quantified in the BALF, and a correlation analysis was performed (Figs. 4D–4F). Peribronchiolitis was more pronounced in the combined exposure group (H-ASD + Con-15H316) than in the Con-15H316 and H-ASD alone groups (Fig. 4A). Similarly, alveolar suppurative inflammation was exacerbated in the combined group relative to the single-agent exposure groups (Fig. 4B). Granulomas, consisting mainly of H-ASD-phagocytosing macrophages, were also more prominent in the combined group compared to the single-agent exposure groups (Fig. 4C). Consistent with these histopathological findings, both LDH and CCL2 levels in the BALF were significantly elevated in the combined group compared to the single-agent exposure groups (Figs. 4D and 4E). Comparing the two single-agent groups, LDH tended to be higher in the Con-15H316 group, while CCL2 levels were higher in the H-ASD group. A negative correlation trend was observed between LDH and CCL2 levels in the single-agent groups (Fig. 4F), implying that these markers reflect distinct inflammatory processes.

**Figure 4.**
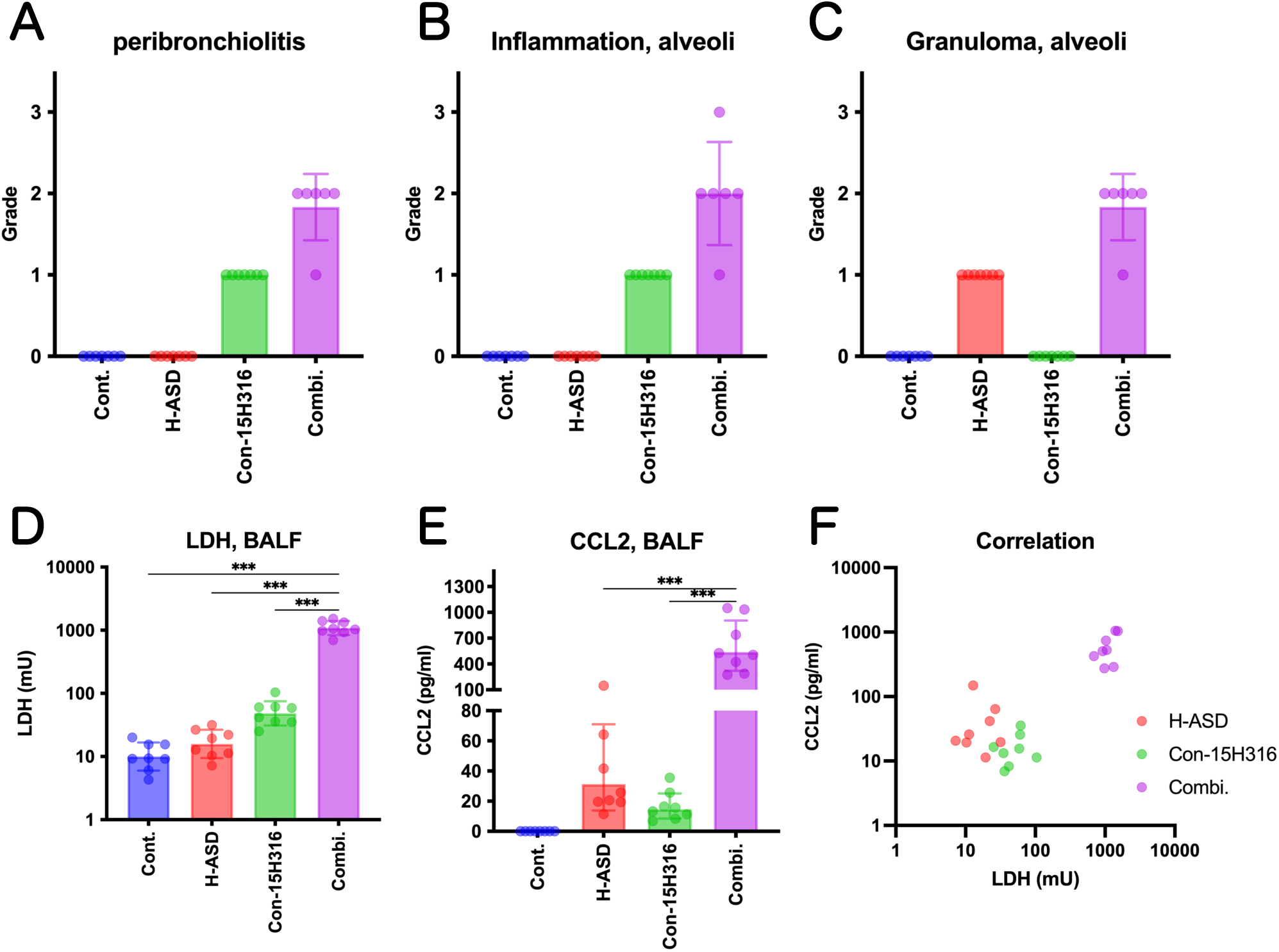
Comparative evaluation of lung lesions and BALF biomarkers. (A) Severity scores of peribronchiolitis across exposure groups. (B) Suppurative inflammation scores in alveolar regions. (C) Granulomatous lesion scores in alveolar regions. (D) Lactate dehydrogenase (LDH) activity in the BALF. (E) CCL2 concentrations in the BALF. (F) Correlation between LDH and CCL2 levels across samples. Bars represent mean ± SD. ***p < 0.001. LDH activity and CCL2 levels were reported in our previous publication (Sadakane *et al*. Int J Environ Health Res 2025) and were re-analyzed and redrawn here for comparative visualization.

Overall, these results demonstrate lesion-specific pathophysiological patterns across exposure groups. Alveolar suppurative inflammation correlated with LDH activity, whereas granulomatous inflammation involving macrophages appeared to correlate with CCL2 levels in the BALF.

### Single-Cell Transcriptomic Analysis Reveals Myeloid Lineage Cells as the Primary Source of CCL2 in Microbial Lung Injury

To clarify the cellular origin of CCL2 (MCP-1) observed in the BALF, we conducted a reanalysis of publicly available single-cell RNA-seq datasets from models of microbial lung injury induced by biologically relevant airborne microorganisms analogous to the fungal exposure in our study. Unsupervised clustering and UMAP projections revealed diverse immune and structural cell populations (Figs. 5A and 5C). In both datasets, Ccl2 expression was highest in clusters annotated as monocytes and interstitial macrophages (IMs). These Ccl2+ populations frequently co-expressed Adgre1 and C1qa–c, hallmark genes of activated monocyte/macrophage lineages (Figs. 5B and 5D). In contrast, epithelial (Epcam+) cells exhibited negligible Ccl2 expression.

**Figure 5.**
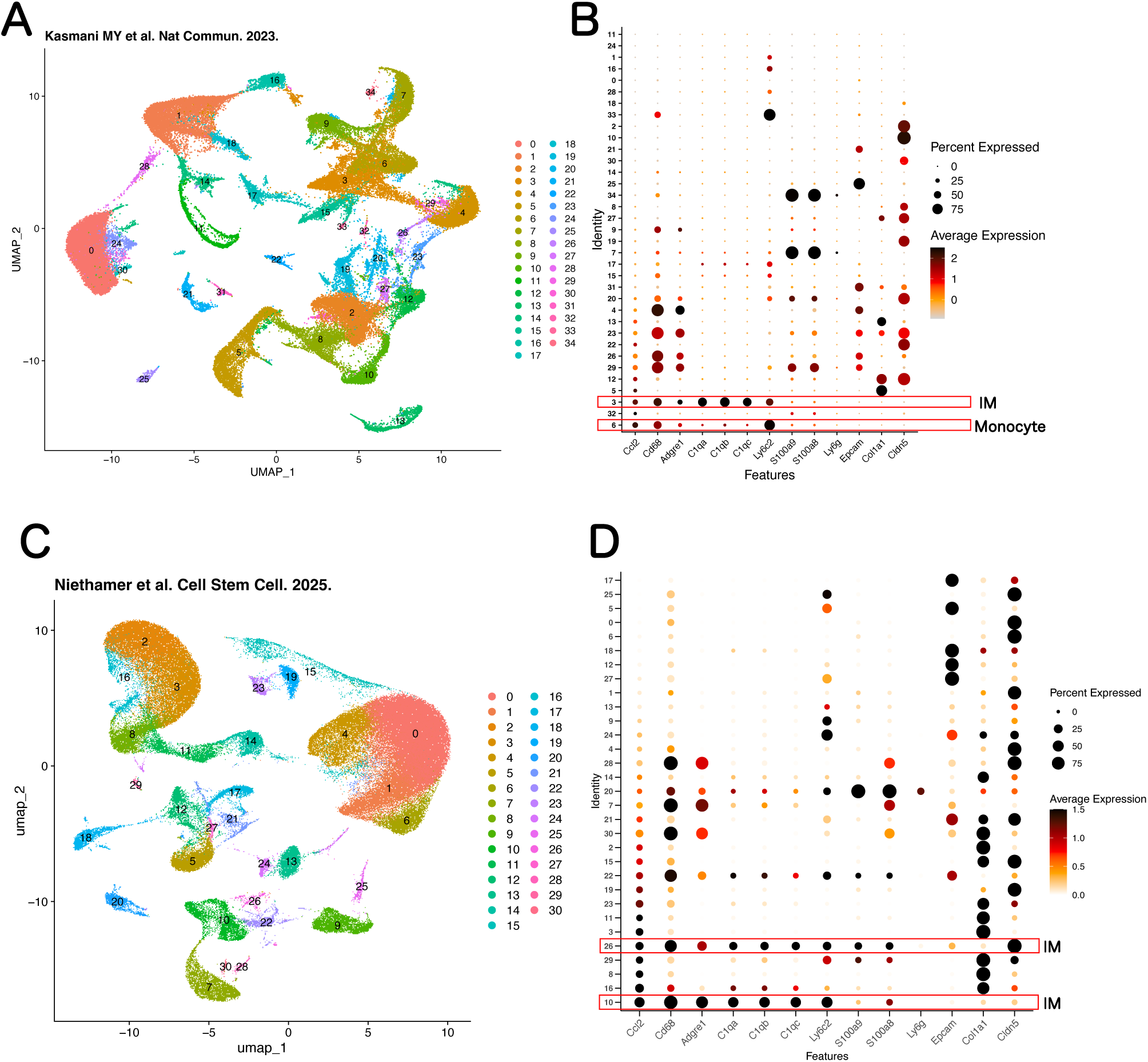
Identification of Ccl2 (MCP-1)–expressing cell populations using single-cell RNA-seq. (A, C) UMAP projections of lung cells from influenza-infected mice (Niethamer et al., 2025; Kasmani et al., 2023) reveal diverse immune and structural cell types. (B, D) Dot plots show highest Ccl2 expression in CD68+ monocytes and interstitial macrophages, co-expressing Ly6c2, S100a8/9, and C1qa–c.

Consequently, the BALF CCL2 elevation observed in our murine exposure model is likely attributable to activated myeloid-lineage cells recruited to the alveolar space in response to H-ASD and/or fungal components. This provides a cellular basis linking cytokine measurements in the BALF to transcriptional activation at single-cell resolution.

## Discussion

Heterogeneous ASD aerosols containing mineral and fungal particles are known to exacerbate the risk of respiratory disorders, as biological and non-biological components trigger different pathways associated with allergic reactions to induce synchronous immune responses ^10,23,24^. Consequently, ASD can trigger respiratory disorders in people without respiratory allergies as well as exacerbate respiratory disorders in people with respiratory allergies. To clarify the mechanisms of the synchronous respiratory effects, histopathological alterations and inflammatory profiles were analyzed using mouse lungs exposed to ASD mineral particles (H-ASD) and the associated fungus (*C. fuckelii*: Con-15H316). Distinct lesion patterns corresponded to the exposure type. H-ASD alone induced granulomas characterized by macrophage-dominant micronodules in alveolar spaces. Con-15H316 alone caused peribronchiolitis rich in eosinophils and lymphocytes. Co-exposure to both agents resulted in induction of granulomatous and suppurative alveolar lesions in combination with airway inflammation and goblet cell hyperplasia, resulting in more widespread and severe inflammation involving both bronchial and alveolar compartments and suggesting additive or synergistic pathogenic mechanisms (Fig. 6). Much of our previous studies primarily focused on allergic airway inflammation and cytokine responses under similar exposure conditions^10,23–27^. The re-analyses of archived lung samples and BALF data adds to our previous findings and highlights a role for suppurative lesions that has not been previously characterized.

**Figure 6.**
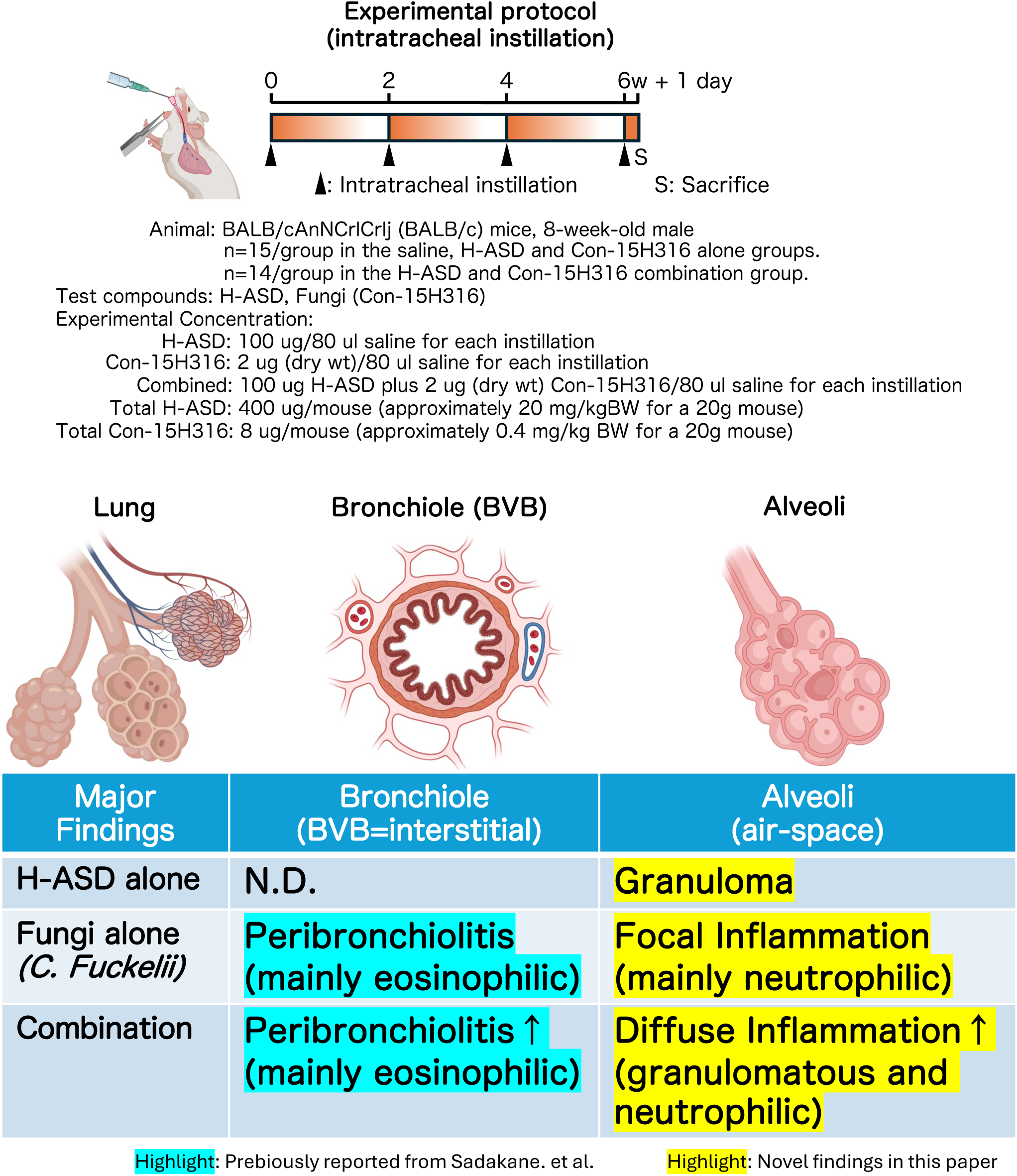
Graphical abstract summarizing key pathological findings. Schematic representation of experimental findings across lung compartments. H-ASD alone induced macrophage-dominant granulomas in alveolar regions, while Con-15H316 provoked eosinophilic peribronchiolitis. Notably, co-exposure resulted in a distinct and synergistically aggravated phenotype, combining granulomatous and suppurative alveolar lesions with airway inflammation and goblet cell hyperplasia. This figure was partially created with BioRender.com.

*C. fuckelii* is a dematiaceous fungus known primarily as a plant pathogen, particularly associated with rose stem canker and dieback in woody ornamental plants. Its pathogenesis has been documented in various horticultural settings through invasion of damaged plant tissues, degradation of periderm layers, and the formation of characteristic pycnidia ^28–30^. To date, there are no published reports confirming its environmental presence in Japan, suggesting that the fungal population (Con-15H316) likely originated from continental East Asia and was transported to Japan via long-range atmospheric transport, possibly during dust storm events. This finding suggests the potential for airborne plant pathogens to cross marine boundaries and seed downwind ecosystems—including mammalian respiratory systems. This study is the first to demonstrate that in addition to plant disease *C. fuckelii*, and likely other fungus species, has the ability to induce lung pathology in mammals.

Previous studies, including our earlier work by Sadakane et al.^25–27^, emphasized allergic responses such as eosinophilic inflammation and cytokine elevation in asthma-like models involving microbial elements (e.g., β-glucan and lipopolysaccharide) and ASD. In contrast, the current reanalysis uncovered a qualitatively distinct pathology: severe non-allergic, suppurative pneumonia in the alveolar regions of mice co-exposed to H-ASD and Con-15H316. This type of inflammation—characterized by neutrophilic infiltration, cellular debris, and macrophage activation—was not prominent in single-agent exposures. Its emergence under co-exposure conditions highlights a previously underappreciated risk of airborne fungi: the induction of innate, dominant alveolar injury in addition to airway inflammation affiliated with allergy associated pathways. While the respiratory toxicity of ASD has been broadly evaluated using cytokine profiling and immune cell quantification, few studies have addressed the lesion-specific distribution and morphology of inflammation at the tissue level. Our study fills this gap by providing diagnostic, visual, and topographical evidence that different lung compartments (airway vs. alveolar) respond uniquely to the combination of mineral and fungal particles.

We have previously reported that systemic inhalation exposure to titanium dioxide nanoparticles (TiO₂ NP) induces interstitial lung disease in rats^17,19^, but not in mice^16^, under OECD guideline-compliant conditions. In this study, H-ASD-laden macrophages were exclusively observed in alveolar spaces, not in bronchovascular bundles or vascular-associated interstitium of mouse lungs. This compartmental restriction of granulomas suggests that mice may be relatively resistant to certain forms of interstitial fibrosis. Thus, the employment of alternative models using larger mammals is warranted to more accurately assess the broader risk of bioaerosol-induced pneumoconiosis and chronic lung disease in occupational and environmental settings.

BALF analysis also revealed synergistic increases in both CCL2 (MCP-1) level and lactate dehydrogenase (LDH) activity in the co-exposure group. The chemokine CCL2, which recruits monocytes and promotes macrophage activation^31^, was particularly elevated in the H-ASD and co-exposure groups. The re-analysis using scRNA-seq supported this result, identifying monocytes and interstitial macrophages as the dominant Ccl2-expressing cells in infected murine lungs. This transcriptional signature would elevate CCL2 and recruit innate immune cells to granulomatous lesions. Conversely, LDH, a marker of cell damage, was higher in the Con-15H316 group, consistent with its tissue-destructive potential, particularly in the bronchial epithelium and adjacent interstitium. The inverse relationship between the mineral elevation of CCL2 and the microbial induction of LDH suggests distinct inflammatory axes: chemokine-driven monocyte recruitment versus cytotoxic damage mediated by eosinophils or necrotic cell death. The co-exposure model appears to unite both inflammatory modes, resulting in a hybrid pathology of granulomatous and suppurative lesions, not captured by conventional asthma models^25^. This lung inflammation caused by heterogenous-agent particles closely mirrors actual environmental exposures, where individuals inhale complex combinations of mineral particles, organic matter, and airborne microorganisms. These findings suggests that neutrophil-driven, non-allergic lung injury be added to current risk assessment of respiratory exposure to particulates and bioaerosols, which often focus allergic mechanisms.

Limitations of this study include the use of a single fungal strain and an acute exposure protocol, which may not fully recapitulate the chronic or repeated exposures encountered in real-world environments. Additionally, our findings are based on a murine model, and extrapolation to human disease requires caution. Future studies should address a broader range of fungal taxa, host susceptibility factors such as age, genetic background, preexisting respiratory conditions, and include long-term exposure scenarios.

## Conclusions

In this study, we conducted a detailed histopathological re-evaluation of murine lung tissues exposed to dust mineral particles (H-ASD), the phytopathogenic fungus *C. fuckelii* (Con-15H316), or both. Consequently, mineral particles in Asian dust were found to induce macrophage-dominant granulomas in alveolar regions, while fungal particle exposure resulted in eosinophilic peribronchiolitis. Notably, co-exposure aggravated distinct and synergistically harmful phenotypes in which granulomatous lesions and neutrophil driven suppurative alveolar lesions developed in combination with airway peribronchiolitis.

These findings were paralleled by co-elevations of lactate dehydrogenase (LDH) and CCL2 (MCP-1) in bronchoalveolar lavage fluid (BALF), indicating simultaneous tissue damage and monocyte-driven inflammation. Additionally, the re-analysis by single-cell RNA-seq could identify monocytes and interstitial macrophages as the primary Ccl2-expressing populations under microbial stress.

To our knowledge, this is the first study to demonstrate that *C. fuckelii* — previously known only as a plant pathogen— can contribute to pulmonary inflammation in mammals, particularly in the context of mineral dust co-exposure. Our results reveal a previously unrecognized, non-allergic form of alveolar pneumonia induced by microbial synergy. These insights underscore the need to expand environmental risk assessment frameworks to account for bioaerosol–dust interactions, which may pose underestimated respiratory hazards in actual exposure conditions.

## Material and Methods

### Collection and Preparation of Aerosol and Microbial Samples

Aerosol-particle sampling was performed at altitudes ranging from 400 to 2,500 m over the Noto Peninsula, Japan, using helicopter- and balloon-based sampling^5,10^. Sampling sites covered Uchinada Town, Hakui City, and Suzu City located around Noto Peninsula (36.7–37.5°N, 136.6–137.4°E). Aerosol particles were collected using sterile pore-size 0.22 μm polycarbonate filters (Whatman, UK) mounted in autoclaved filter holders (Swinnex, Merck, Germany) at a flow rate of 5 L/min for 0.2–1.0 h. After sampling, the filters were stored at a temperature of −80 °C until further processing. Fungi were isolated by aerosol suspension in potato dextrose (PD) medium and identified by internal transcribed spacer (ITS) sequencing using SR1R and ITS4 primers^35^. The isolate *C. fuckelii* (Con-15H316) was selected for the present study. Following established protocols^24,33,34^, after fungal hyphae were fixed with 1% formalin for 24 h at 4 °C, the fixed fungal hyphae were washed and sonicated for 1 min on ice using a Microchip Sonicator (UD-201; Tomy Digital Biology Co., Tokyo, Japan). The washing process was repeated five times. Final pellet weights were determined gravimetrically after drying. A standard Asian sand dust (ASD) sample (Gobi Kosa Dust No. 30; National Institute for Environmental Studies, Japan) was sieved to <2.5 μm, sterilized by heating at 360 °C for 30 min in an electric furnace (SSTR-25K; Isuzu CAP, Niigata, Japan), and stored in sterile dry bottles at −30 °C. Particle size distribution was assessed using a fluorescence microscope (BZ-9000; KEYENCE, Osaka, Japan). The median particle diameter was 2.87 ± 2.42 μm, with a peak distribution between 1.0 and 2.0 μm^25,35^.

The collection, preparation, animal exposure, and BALF sampling procedures are described in our previous study^10^. In the present study, we performed additional histopathological re-analysis using newly prepared lung tissue sections.

### Animal Models and Experimental Design

The original experimental design is indicated in Fig. S2. Eight-week-old male BALB/c mice (Charles River Japan) were acclimated for one week and housed under specific pathogen-free (SPF) conditions with ad libitum access to a CE-2 diet (CLEA Japan) and water. The animal facility was maintained at 23–25 °C with 55–70% humidity under a 12-h light/dark cycle.

Mice were randomly assigned to four experimental groups:

1. Saline control: n=8 for BALF, n=7 for histopathology
2. H-ASD alone: n=8 for BALF, n=7 for histopathology
3. C. fuckelii (Con-15H316) alone: n=8 for BALF, n=7 for histopathology
4. H-ASD + Con-15H316: n=8 for BALF, n=6 for histopathology

Each treatment (80 μL) contained 0.1 mg H-ASD, 2 μg of fungal dry weight, or both suspended in sterile, endotoxin- and β-glucan-free saline. Suspensions were sonicated for 5 min on ice (UD-201; Tomy Digital Biology Co., Ltd., Tokyo, Japan). Mice were anesthetized with 4% halothane and administered the suspension intratracheally via polyethylene tubes. Dosing was performed four times at two-week intervals, in accordance with previous protocols^27,34,35^. Mice were euthanized 24 h after the final dose by intraperitoneal injection of pentobarbital.

### Bronchoalveolar Lavage Fluid (BALF) Collection and Analysis

BALF was collected from eight mice per group by intratracheal instillation of 0.8 mL warm (37 °C) sterile saline. BALF collection was repeated twice. The recovered fluid was pooled and centrifuged at 402 × g for 10 min at 4 °C. Supernatants were stored at −80 °C until analysis.

Lactate dehydrogenase (LDH) activity was measured using a commercial LDH Assay Kit (Abcam, Cambridge, UK; Cat# ab102526; Lot# GR3268979-1), with a detection limit of 1 mU/mL. Monocyte chemoattractant protein-1 (MCP-1) concentrations were determined using a Quantikine ELISA Kit for mouse CCL2/MCP-1 (R&D Systems, Minneapolis, MN, USA; Cat# MJE00B; Lot# P307801), with a detection limit of 2 pg/mL. All assays were conducted in duplicate in accordance with manufacturer protocols^10,33^.

In the present study, no new BALF collections were conducted; previously obtained data was re-analyzed for this study.

### Histopathological analysis

Formalin-fixed, paraffin-embedded lung blocks obtained in our previous study^10^ were used. For this study, new thin sections (2-μm thick) were prepared from these archived blocks and stained with hematoxylin and eosin (H&E) for additional histopathological evaluation. Histopathological diagnoses were made independently by two board-certified toxicologic pathologists (Shotaro Yamano and Yumi Umeda) from the Japanese Society of Toxicologic Pathology, following the International Harmonization of Nomenclature and Diagnostic Criteria for Lesions in Rats and Mice (INHAND) ^36^ and the diagnostic reference for human pneumoconiosis^37,38^. Non-neoplastic lesions were graded for severity on a scale from “slight” to “severe” according to the criteria established by Shackelford et al.^39^. Microscopic examination was performed using an ECLIPSE Ni optical microscope (Nikon Corp., Tokyo, Japan), a BZ-X810 (KEYENCE, Osaka, Japan), or the VS120 virtual slide scanner (Olympus, Tokyo, Japan).

### Re-analysis of publicly available single-cell RNA-seq datasets

To better understand the cellular sources of Ccl2 (MCP-1) in the lung following microbial challenge, we reanalyzed publicly available single-cell RNA sequencing (scRNA-seq) datasets generated from mice exposed to infectious agents. This analysis was conducted to provide context for our observed increases in CCL2 levels in the bronchoalveolar lavage fluid (BALF) following intratracheal administration of C. fuckelii and/or H-ASD. Specifically, two datasets were selected: GSE202325 (Niethamer et al., Cell Stem Cell, 2025)^40^ and GSE229678 (Kasmani et al., Nature Communications, 2023)^41^, both derived from murine models of influenza infection. These datasets represent respiratory exposure to biologically relevant microbial stimuli, analogous to the fungal challenge employed in our study.

Raw gene expression matrices were obtained from the Gene Expression Omnibus (GEO) and the Chan Zuckerberg Initiative’s cellxgene platform (https://cellxgene.cziscience.com/). The GSE202325 dataset, provided in .h5ad format, was converted and imported into R using SeuratDisk for compatibility with Seurat. Data processing was performed using R version 4.5.1 (2025-06-13) and Seurat version 5.3.0. After filtering out cells with fewer than 200 detected genes or over 15% mitochondrial content, each dataset was log-normalized and integrated using the canonical correlation analysis (CCA) pipeline. Dimensionality reduction was achieved using principal component analysis (PCA) followed by Uniform Manifold Approximation and Projection (UMAP), and clustering was performed using the Louvain algorithm. We specifically investigated the cell types responsible for Ccl2 expression by visualizing gene-level expression patterns across major immune and structural cell populations. UMAP and dot plots were generated using Seurat’s DimPlot() and DotPlot() functions.

Marker genes included Cd68, Adgre1, C1qa, C1qb, C1qc, Ly6c2, S100a8, S100a9, and Ly6g for macrophages, monocytes, and neutrophils; Epcam and Cldn5 for epithelial and endothelial cells; Col1a1 for fibroblasts; and Ccl2 for chemokine signaling. All visualizations were exported as high-resolution vector graphics (300 dpi) using ggsave(). To aid interpretation, dot plots were manually reordered based on Ccl2 expression intensity. This reanalysis enabled identification of putative Ccl2-producing cell populations and provided insight into the potential cellular contributors to the elevated BALF CCL2 levels observed in our murine exposure model.

## Statistical analysis

For histopathological lesion scores, comparisons among multiple groups were performed using the Kruskal–Wallis test, followed by Dunn’s multiple comparisons test when significant differences were detected. For bronchoalveolar lavage fluid (BALF) biomarkers, including lactate dehydrogenase (LDH) activity and CCL2 (MCP-1) levels, data were analyzed using one-way ANOVA. When the ANOVA was significant, Tukey’s multiple comparison test was used to assess differences between groups. All statistical analyses were conducted using GraphPad Prism software (version 10.5.0; GraphPad Software, San Diego, CA, USA). Statistical significance was defined as p < 0.05.

## Abbreviations

ASD: Asian Sand Dust
BALF: Bronchoalveolar lavage fluid
Ccl2: Chemokine (C-C motif) ligand 2
CWAAP: Cross-linked water-soluble acrylic acid polymer
ELISA: Enzyme-linked immunosorbent assay
GEO: Gene Expression Omnibus
H-ASD: Heat-treated Asian Sand Dust
H&E: Hematoxylin and eosin
IM: Interstitial macrophage
ITS: Internal transcribed spacer
LDH: Lactate dehydrogenase
LPS: Lipopolysaccharide
MCP-1: Monocyte chemoattractant protein-1
MWCNT: Multi-walled carbon nanotube
OECD: Organisation for Economic Co-operation and Development
PCA: Principal component analysis
scRNAseq: Single-cell RNA sequencing
SPF: Specific pathogen-free
TiO₂ NP: Titanium dioxide nanoparticle
UMAP: Uniform manifold approximation and projection

## Declarations

### Ethics approval and consent to participate

This study was performed in accordance with the Animal Research: Reporting of In Vivo Experiments (ARRIVE) guidelines and the U.S. National Institutes of Health Guidelines for the Care and Use of Laboratory Animals. The animal care method was approved by the Animal Care and Use Committee of the Oita University of Nursing and Health Sciences, Oita, Japan (approval number: 17–23). The lung tissue sections used in this study were newly prepared from archived paraffin-embedded blocks obtained in our previous approved study^10^.

### Consent for publication

All authors gave their consent for publication of this manuscript.

### Availability of data and materials

The datasets used and analyzed during the current study are available from the corresponding authors on reasonable request.

## Acknowledgments

We appreciate the vital contributions of the students at the Oita University of Nursing and Health Sciences to this research. We wish to thank Dr. David B. Alexander of Nanotoxicology project, Nagoya City University Graduate School of Medicine for his insightful comments and English editing. We would also like to thank Misae Saito for her assistance with the pathological experiments.

## Author information

National Institute of Occupational Safety and Health, Japan Organization of Occupational Health and Safety, Fujisawa, Kanagawa 251-0015, Japan Shotaro Yamano, Yumi Umeda

Department of Health Sciences, Oita University of Nursing and Health Sciences, Oita, Oita 870-1201, Japan

Kaori Sadakane, Takamichi Ichinose

Graduate School of Global Environmental Studies, Kyoto University, Kyoto, 615-8246, Japan

Takamichi Ichinose

Department of Life Science, Faculty of Science and Engineering, Kindai University, Higashiosaka, Osaka 577-8502, Japan

Teruya Maki

## Contributions

All authors contributed to the conception and design of this study. Material preparation, data collection, and analyses were performed by T.M, S.Y, K.S, T.I, and Y.U. The first draft of the manuscript was written by S.Y, and all authors commented on previous versions of the manuscript. All the authors have read and approved the final version of the manuscript.

## Supplementary Information

Additional file 1

**Figure S1.**
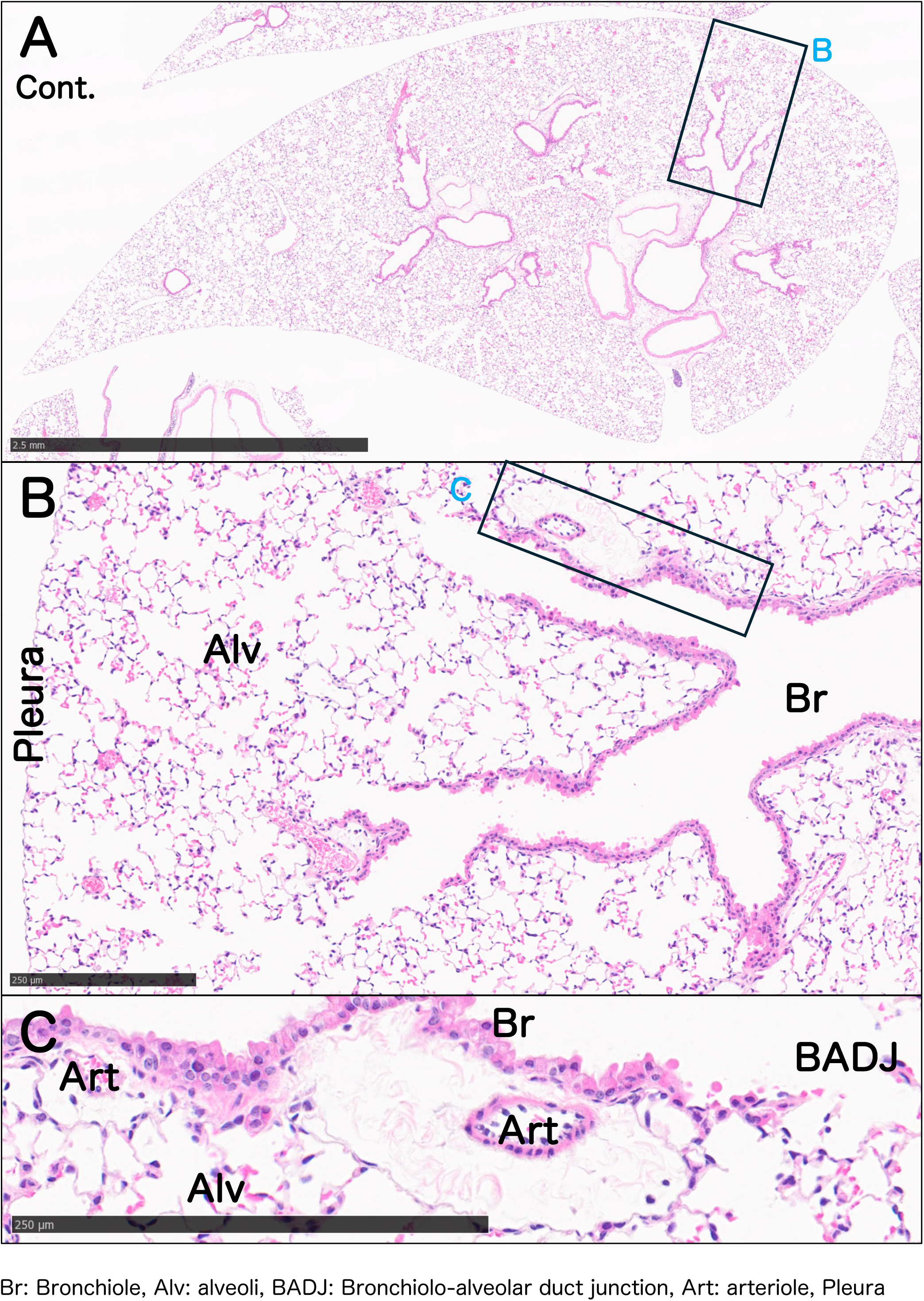
Histological characteristics of normal mouse lung histology. Representative H&E images of control lung sections. Bronchioles (Br), alveoli (Alv), bronchiole–alveolar duct junctions (BADJ), arterioles (Art), and pleura are labeled. Normal histoarchitecture is preserved, providing a reference for lesion comparison.

Additional file 2

**Figure S2.**
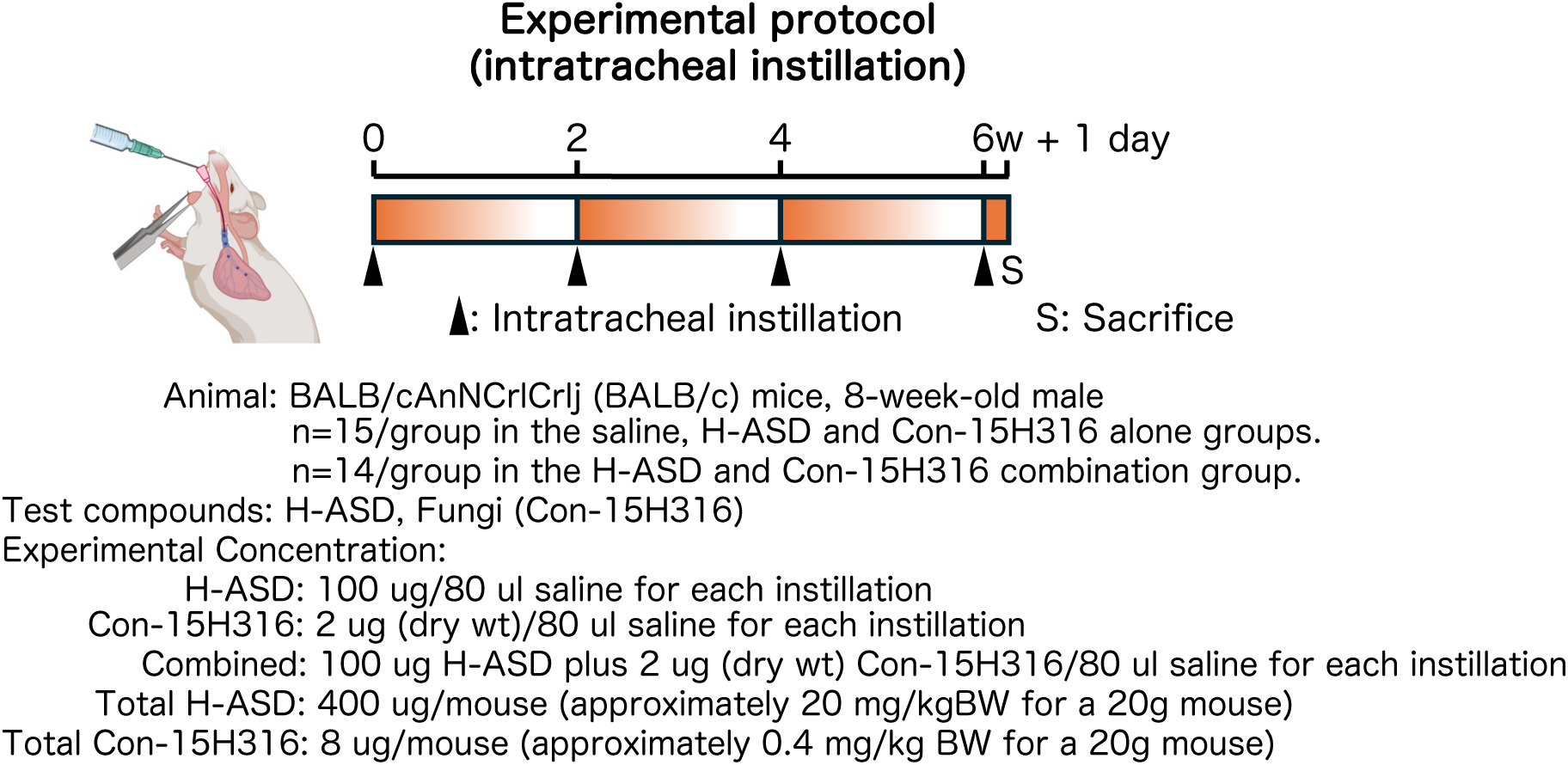
Experimental design schematic. BALB/c mice (n=15/each group in saline, H-ASD and Con-15H316 alone, and n=14/group in combination) were intratracheally administered saline, H-ASD, Con-15H316, or a combination of H-ASD and Con-15H316 on days 0, 14, 28, and 42. Sacrifice and tissue collection was carried out 24 h after the final dose (day 43). Total doses: H-ASD, 400 µg/mouse; Con-15H316, 8 µg/mouse. Key endpoints: histopathology and BALF biomarker analysis.

## References

1. Fussell, J. C. & Kelly, F. J. Mechanisms underlying the health effects of desert sand dust. Environ Int 157, 106790 (2021).

2. Duce, R. A., Unni, C. K., Ray, B. J., Prospero, J. M. & Merrill, J. T. Long-Range Atmospheric Transport of Soil Dust from Asia to the Tropical North Pacific: Temporal Variability. Science 209, 1522–1524 (1980).

3. Cao, C. et al. Inhalable Microorganisms in Beijing’s PM2.5 and PM10 Pollutants during a Severe Smog Event. Environ. Sci. Technol. 48, 1499–1507 (2014).

4. Yeo, H.-G. & Kim, J.-H. SPM and fungal spores in the ambient air of west Korea during the Asian dust (Yellow sand) period. Atmospheric Environment 36, 5437– 5442 (2002).

5. Maki, T. et al. Variations in airborne bacterial communities at high altitudes over the Noto Peninsula (Japan) in response to Asian dust events. Atmospheric Chemistry and Physics 17, 11877–11897 (2017).

6. Kwon, H.-J., Cho, S.-H., Chun, Y., Lagarde, F. & Pershagen, G. Effects of the Asian Dust Events on Daily Mortality in Seoul, Korea. Environmental Research 90, 1–5 (2002).

7. Yang, C.-Y., Tsai, S.-S., Chang, C.-C. & Ho, S.-C. Effects of Asian dust storm events on daily admissions for asthma in Taipei, Taiwan. Inhal Toxicol 17, 817– 821 (2005).

8. Park, J. W. et al. Effects of ambient particulate matter on peak expiratory flow rates and respiratory symptoms of asthmatics during Asian dust periods in Korea. Respirology 10, 470–476 (2005).

9. Watanabe, M. et al. Correlation between Asian Dust Storms Worsening Asthma in Western Japan. Allergology International 60, 267–275 (2011).

10. Sadakane, K., Ichinose, T., Maki, T., Takano, H. & Shibamoto, T. A comparative analysis of five fungal species isolated from high-altitude air samples based on their induction of murine lung eosinophilia along with heated Asian sand dust. Int J Environ Health Res 1–15 (2025) doi:10.1080/09603123.2025.2484775.

11. OECD. Test No. 413: Subchronic Inhalation Toxicity: 90-Day Study. (Organisation for Economic Co-operation and Development, Paris, 2018).

12. OECD. Test Guideline No. 451: Carcinogenicity Studies. (Organisation for Economic Co-operation and Development, Paris, 2018).

13. Umeda, Y. et al. Two-week Toxicity of Multi-walled Carbon Nanotubes by Whole-body Inhalation Exposure in Rats. J Toxicol Pathol 26, 131–140 (2013).

14. Kasai, T. et al. Thirteen-week study of toxicity of fiber-like multi-walled carbon nanotubes with whole-body inhalation exposure in rats. Nanotoxicology 9, 413– 422 (2015).

15. Kasai, T. et al. Lung carcinogenicity of inhaled multi-walled carbon nanotube in rats. Part Fibre Toxicol 13, 53 (2016).

16. Yamano, S. et al. No evidence for carcinogenicity of titanium dioxide nanoparticles in 26-week inhalation study in rasH2 mouse model. Sci Rep 12, 14969 (2022).

17. Yamano, S. et al. Pulmonary dust foci as rat pneumoconiosis lesion induced by titanium dioxide nanoparticles in 13-week inhalation study. Part Fibre Toxicol 19, 58 (2022).

18. Kasai, T. et al. Lung carcinogenicity by whole body inhalation exposure to Anatase-type Nano-titanium Dioxide in rats. J Toxicol Sci 49, 359–383 (2024).

19. Yamano, S. & Umeda, Y. Fibrotic pulmonary dust foci is an advanced pneumoconiosis lesion in rats induced by titanium dioxide nanoparticles in a 2-year inhalation study. Particle and Fibre Toxicology 22, 7 (2025).

20. Takeda, T. et al. Dose–response relationship of pulmonary disorders by inhalation exposure to cross-linked water-soluble acrylic acid polymers in F344 rats. Particle and Fibre Toxicology 19, 27 (2022).

21. Yamano, S. et al. Mechanisms of pulmonary disease in F344 rats after workplace-relevant inhalation exposure to cross-linked water-soluble acrylic acid polymers. Respir Res 24, 47 (2023).

22. Goto, Y. et al. Carcinogenicity and testicular toxicity of 2-bromopropane in a 26-week inhalation study using the rasH2 mouse model. Sci Rep 13, 1782 (2023).

23. He, M. et al. Asian sand dust enhances murine lung inflammation caused by Klebsiella pneumoniae. Toxicol Appl Pharmacol 258, 237–247 (2012).

24. Liu, B. et al. Lung inflammation by fungus, Bjerkandera adusta isolated from Asian sand dust (ASD) aerosol and enhancement of ovalbumin-induced lung eosinophilia by ASD and the fungus in mice. Allergy Asthma Clin Immunol 10, 10 (2014).

25. Sadakane, K., Ichinose, T., Nishikawa, M., Takano, H. & Shibamoto, T. Co-exposure to zymosan A and heat-inactivated Asian sand dust exacerbates ovalbumin-induced murine lung eosinophilia. Allergy Asthma Clin Immunol 12, 48 (2016).

26. Sadakane, K., Ichinose, T. & Nishikawa, M. Effects of co-exposure of lipopolysaccharide and β-glucan (Zymosan A) in exacerbating murine allergic asthma associated with Asian sand dust. J Appl Toxicol 39, 672–684 (2019).

27. Sadakane, K., Ichinose, T., Maki, T. & Nishikawa, M. Co-exposure of peptidoglycan and heat-inactivated Asian sand dust exacerbates ovalbumin-induced allergic airway inflammation in mice. Inhal Toxicol 34, 231–243 (2022).

28. Alfieri, S. A. Jr. STEM AND GRAFT CANKER OF ROSE. Plant Pathology Circular 88, (1969).

29. Jamali, S. First report of *Paraconiothyrium fuckelii* (*Didymosphaeriaceae*, Pleosporales), causing stem canker of Rosa hybrida, from Iran. Czech Mycol. 72, 71–82 (2020).

30. Zaher, E., Attia, M. & Reyad, N. E.-H. Rose Stem Canker Caused by Coniothyrium fuckelii Sacc. in Egypt. Egyptian Journal of Phytopathology 40, 39 (2012).

31. Deshmane, S. L., Kremlev, S., Amini, S. & Sawaya, B. E. Monocyte chemoattractant protein-1 (MCP-1): an overview. J Interferon Cytokine Res 29, 313–326 (2009).

32. Toju, H., Tanabe, A. S., Yamamoto, S. & Sato, H. High-Coverage ITS Primers for the DNA-Based Identification of Ascomycetes and Basidiomycetes in Environmental Samples. PLOS ONE 7, e40863 (2012).

33. He, M. et al. The Role of Toll-Like Receptors and Myeloid Differentiation Factor 88 in Bjerkandera adusta-Induced Lung Inflammation. Int Arch Allergy Immunol 168, 96–106 (2015).

34. He, M. et al. Desert dust induces TLR signaling to trigger Th2-dominant lung allergic inflammation via a MyD88-dependent signaling pathway. Toxicol Appl Pharmacol 296, 61–72 (2016).

35. He, M. et al. Effects of two Asian sand dusts transported from the dust source regions of Inner Mongolia and northeast China on murine lung eosinophilia. Toxicology and Applied Pharmacology 272, 647–655 (2013).

36. Renne, R. et al. Proliferative and nonproliferative lesions of the rat and mouse respiratory tract. Toxicol Pathol 37, 5S–73S (2009).

37. Anna-Luise A. Katzenstein. Diagnostic Atlas of Non-Neoplastic Lung Disease: A Practical Guide for Surgical Pathologists. (Demos Medical, 2016).

38. Mukhopadhyay, S. Non-Neoplastic Pulmonary Pathology with Online Resource: An Algorithmic Approach to Histologic Findings in the Lung. (Cambridge University Press, Cambridge, 2016).

39. Shackelford, C., Long, G., Wolf, J., Okerberg, C. & Herbert, R. Qualitative and quantitative analysis of nonneoplastic lesions in toxicology studies. Toxicol Pathol 30, 93–96 (2002).

40. Niethamer, T. K. et al. Longitudinal single-cell profiles of lung regeneration after viral infection reveal persistent injury-associated cell states. Cell Stem Cell 32, 302–321.e6 (2025).

41. Kasmani, M. Y. et al. A spatial sequencing atlas of age-induced changes in the lung during influenza infection. Nat Commun 14, 6597 (2023).

